# Disease decreases variation in host community structure in an old-field grassland

**DOI:** 10.1101/2022.08.15.503989

**Authors:** Rita L. Grunberg, Fletcher W. Halliday, Robert W. Heckman, Brooklynn N. Joyner, Kayleigh R. O’Keeffe, Charles E. Mitchell

## Abstract

Disease may modulate variation in host community structure by modifying the interplay of deterministic and stochastic processes. For instance, deterministic processes like ecological selection can benefit species less impacted by disease. When disease consistently selects for certain host species, this can reduce variation in host community composition. On the other hand, when host communities are less impacted by disease and selection is weaker, stochastic processes (e.g., drift, dispersal) may play a bigger role in host community structure, which can increase variation in structure among communities. While effects of disease on host community structure have been quantified in field experiments, few have addressed the role of disease in modulating variation in structure among host communities. To address this, we conducted a field experiment spanning three years, using a tractable system: foliar fungal pathogens in an old-field grassland community dominated by the grass *Lolium arundinaceum*, tall fescue. We reduced foliar fungal disease burden in replicate host communities (experimental plots in intact vegetation) in three fungicide regimens that varied in the duration of fungicide exposure and included a fungicide-free control. We measured host diversity, biomass, and variation in community structure among replicate communities. Disease reduction generally decreased plant richness and increased aboveground biomass relative to communities experiencing ambient levels of disease. Despite changes in structure of the plant communities over the experiment’s three years, the effects of disease reduction on plant richness and biomass were consistent across years. However, disease reduction did not reduce variation in host community structure, providing little evidence for ecological selection by competition or other deterministic processes. Instead, disease reduction tended to amplify variation in host community structure among replicate communities (i.e., within fungicide treatment groups), suggesting that disease diminished the degree to which host communities were structured by stochastic processes. These results of experimental disease reduction both highlight the potential importance of stochastic processes in plant communities and reveal the potential for disease to regulate variation in host community structure.

## Introduction

Disease can be an agent of ecological selection (sensu Vellend 2010) and thereby influence the structure of host communities (Minchella and Scott 1991, Hatcher et al. 2006, Mordecai 2011, Friesen et al. 2020). Field experiments reducing disease in communities have revealed that disease can not only increase local species richness, but also can shift community composition (Allan et al. 2010, Heckman et al. 2016, 2017, Wilfahrt et al. 2020). Relative to such effects of disease on local host community structure, disease impacts on variation in structure among host communities and over time have been investigated less (but see Heckman et al. 2016, 2017, Cappelli et al. 2020, Szefer et al. 2020, Wilfahrt et al. 2020). Thus, it remains unresolved to what degree disease plays a deterministic role in host community assembly. Quantifying these disease impacts on host communities over time is important for understanding how disease interacts with other processes that may drive host community assembly.

Communities are structured both by deterministic and stochastic processes (Chase 2007, Vellend 2010, Germain et al. 2013). To understand how disease affects host community composition, it may be important to explore how disease alters the relative importance of deterministic and stochastic processes. In host communities in which interspecific competition is an agent of ecological selection, disease can decrease the degree to which the community is dominated by better competitors and shift the relative abundance of certain host taxa (Hatcher et al. 2006, Mordecai 2011). However, variation among host communities in species composition can lead to variation among communities in selection by natural enemies, impeding host communities from converging towards a consistent set of species (Lind et al. 2013). Ecological selection, such as by disease, can also reduce the relative influence of stochastic demographic processes, such as dispersal or ecological drift, on community structure (Chase et al. 2009, Vellend 2010). Over time or across multiple local communities, stochastic processes can increase variation in community structure (Germain et al. 2013, Gilbert and Levine 2017). Thus, to the degree that disease reduces the relative influence of stochastic processes within host communities, disease may reduce variation in community structure over time and among communities. Consequently, disease impacts on host communities may depend on the interplay of multiple processes, both deterministic and stochastic, that drive community structure. More broadly, disease may increase, have no effect on, or decrease variation in host community structure, depending on the relative importance of disease, along with other agents of ecological selection, and stochastic processes that structure communities.

The role of disease in structuring host communities may also depend on the structure of the parasite community. Hosts are commonly infected with a diverse community of parasite species that vary in their effects on host fitness and growth (Friesen et al. 2017), so variation in parasite community structure can drive disease impacts on host individuals (Mordecai et al. 2015) and communities (Cappelli et al. 2020, Friesen et al. 2021). Moreover, parasite abundance and diversity both vary temporally. Specifically, infections are often seasonal, which will generate differences in disease burdens over time and contribute to variation in parasite community structure (Sapp and Esch 1994, Grunberg and Sukhdeo 2017). Previous field experiments have found that parasite impacts on the host community can depend on parasite community composition (Cappelli et al. 2020, Friesen et al. 2021) and that parasite community composition can shift seasonally (Sapp and Esch 1994, Grunberg and Sukhdeo 2017, Halliday et al. 2017). But, the effect of seasonal variation in parasite community composition on plant community structure remains untested.

Here, we manipulated foliar fungal disease burden by treating plant communities with fungicide for different periods of the growing season, and then tested the effects of foliar fungal disease on changes in plant community structure and biomass over three years. At the beginning of our experiment, plant communities were dominated by the perennial grass tall fescue, which is host to a diverse community of foliar fungal diseases that vary in infection prevalence over the growing season (Halliday et al. 2017). Our seasonal fungicide treatments were designed to alter the species composition of the parasites that cause these different diseases. The fungicide treatments ended up having little effect on parasite species composition (unpublished data), but substantially reduced overall disease burden, so here we focus on the effects of disease burden on host communities. To investigate the interplay of deterministic and stochastic processes, we quantified not only effects of disease on the species richness, composition, and biomass of local host communities, but also their impacts on variation among communities over time.

## Methods

### Experimental design

Our experiment was conducted at Widener Farm in the Duke Forest Teaching and Research Laboratory in Orange Co., North Carolina, USA. This old field site was previously used to produce crops until 1992. Since then, the site has been mowed at least once a year in the summer to maintain dominance of the plant community by herbaceous plants. Typically, four non-native perennial species, *Lespedeza cuneata, Lonicera japonica, Sorghum halepense*, and *Lolium arundinaceum* (= *Festuca arundinacea*, tall fescue) constitute the majority of community cover (Heckman et al. 2016). Of these plant species, tall fescue was the most abundant in our plots (Figure S1) and supports both a heavy load of foliar fungal disease and a great diversity of foliar fungal parasites (Halliday et al. 2019). So, we used observed epidemics of foliar fungal disease on the abundant host tall fescue to design our experimental treatments to shift parasite community composition on that host.

Epidemics of foliar fungal disease on tall fescue are seasonal; typically, the disease anthracnose (caused chiefly although not exclusively by *Colletotrichum cereale)* peaks in late spring, while brown patch (caused by *Rhizoctonia solani*) peaks in the summer and crown rust (caused by *Puccinia coronata)* peaks in the early fall (Figure S2, Halliday et al. 2017). Hereafter, we refer to these diseases rather than the causal parasite species because our study focuses on disease impacts. We designed different fungicide treatment regimens to correspond with the peak of the seasonal epidemics of these diseases, so that different fungicide treatments would shift the composition of disease infecting tall fescue. Treatments varied in their duration of fungicide exposure including: no fungicide (control), fungicide until mid-July (approximately 7 months, soon after the typical start of brown patch epidemics), fungicide until mid-September (approximately nine months, soon after the typical start of crown rust epidemics), and year-round application of fungicide. Fungicide application started each year in January, except in 2017, when fungicide treatments started in May. Throughout, we refer to these fungicide treatments with respect to their designated yearly duration of fungicide application: never, seven months, nine months, and year-round. Fungicide treatments started in May 2017 and ended in February 2020.

In total, we established 64 experimental plots in an intact old-field vegetation site that was fenced to exclude vertebrates, particularly deer and rodents. At the beginning of the experiment on 11-13 June 2017, we increased tall fescue dominance of plant communities by clipping shoots of *Lespedeza cuneata* and *Sorghum halepense* at their base. Experimental plots were assigned to one of four fungicide treatments in a fully randomized design. Each plot was 2 m × 2 m and separated by 1 m with a 2-m buffer surrounding the entire experimental area, and the 64 plots were arranged in a 16×4-plot array with a total footprint of 51 m × 15 m.

The fungicide *Dithane75DF Rainshield*, (75% mancozeb, Dow AgroSciences, Indianapolis, Indiana, USA) was used to reduce foliar fungal disease. Prior studies have found that application of this fungicide at recommended rates did not affect plant growth or mycorrhizal colonization (Parker and Gilbert 2007), including growth of tall fescue and other common species from our experimental site (Heckman et al. 2016). Fungicide was applied approximately every two weeks during the designated fungicide treatment time. Plots not assigned to receive fungicide treatments were instead sprayed with water. In 2018, plots were washed with water at the end of each seasonal treatment’s fungicide application period (i.e., in mid-July for the seven-month treatment and mid-September for the nine-month treatment) to remove fungicide from leaves and allow fungal colonization of the plots for the rest of the year. Plots in all treatments were washed for the same duration to equalize water addition among plots. Plots were mown annually at the end of each growing season to reduce the establishment of woody vegetation, which tends to outcompete herbaceous species in Eastern North American old-fields, including our site (Wright and Fridley 2010, Heckman et al. 2022)

### Data collection

Plant community composition was recorded on 09 November 2017, 10 October 2018, and 10 October 2019. The cover of all plant species, including litter and bare ground, was visually estimated within a 0.75 m x 0.75 m permanent quadrat in each plot. The absolute cover of each plant species was estimated independently as plants may overlap, and thus total cover often exceeded 100% in plots. We then calculated relative cover for each species in a plot as the absolute cover of that species divided by the sum of absolute cover of all species in a plot.

We evaluated the effectiveness of the fungicide treatments by surveying disease prevalence in tall fescue. Disease prevalence was not assayed for other plant species; we rationalized that quantifying disease on the most abundant host species, tall fescue, which initially constituted ∼87% of plant cover in our plots (Figure S1.B), provides a useful estimate for the overall effectiveness of the fungicide on most diseases in this system. We surveyed disease monthly, starting in March 2017 and ending in December 2019, by haphazardly selecting 20 tillers (i.e., individual grass shoots) of tall fescue within each plot and recording visible disease symptoms on all leaves on a tiller. We then quantified disease prevalence within a plot as the total number of infected leaves (infected = any amount of disease) divided by all leaves surveyed across the 20 plants, treating leaves as host individuals because fungal infections are localized within a leaf (Halliday et al. 2017).

We measured aboveground plant biomass each year in mid-November by harvesting the entire 0.75 m x 0.75 m quadrat in 2017, and by harvesting two 0.5 m x 0.2 m strips of vegetation in 2018 and 2019. Dead vegetation from the growing season (litter), which did not include prior year’s biomass, was included in biomass measurements. In 2018, we sorted plant biomass into five categories: tall fescue, non-fescue monocots, non-woody dicots, woody dicots, and litter. Aboveground biomass was then oven-dried at 65 °C for at least 72 hours and then weighed.

### Analysis

#### Fungicide treatment effects on disease

We quantified host population-level disease burden by calculating the annual area under the disease progress stairs (AUDPS) (Simko and Piepho 2012) using the monthly disease prevalence survey data. This measure of disease burden (i.e., AUDPS) provides an advantage over other measures like the area under the disease progress curve because it gives a better estimate of the contribution of the first and last observation (Simko and Piepho 2012). To calculate disease burden, we used disease survey data from May until November of each year to be consistent across years; also, this timespan represents the bulk of the growing season in our system. The AUDPS, our measurement of disease burden, was estimated using the ‘agricolae’ R package (Mendiburu 2021). We calculated a cross-disease disease burden that represents total infection pooled across the three most prevalent diseases of tall fescue (anthracnose, brown patch, and crown rust) as an indicator of overall disease burden. We also include disease burden data for each of the three diseases infecting tall fescue. As a measure of parasite composition, we also quantified the relative contribution of each disease’s AUDPS to the cross-species AUDPS. Quantifying the relative contribution of each disease allows us to account for overall differences in disease due to fungicide.

The effects of the fungicide treatments on disease burden over time were evaluated using a repeated measures analysis of variance (ANOVA). In this analysis, we used the annual cross-disease burden (i.e., AUPDS). In the ANOVA we included fungicide treatment, year, and treatment*year interaction as fixed effects, and experimental plot as a random effect. As a measure of disease reduction, we also report the disease burden log response ratio relative to the control-no fungicide plots.

#### Plant diversity and biomass

We used repeated-measures ANOVAs to test the effects of fungicide treatment over time on plant biomass and plant diversity metrics. Plant diversity was quantified using three metrics based on Hill’s series of diversity (Hill 1973) in the ‘vegan’ package (Oksanen et al. 2008). The value of *q* in Hill’s series is related to differences in the weighting of the relative abundance of taxa: taxonomic richness (*q* = 0, no abundance weighting), Hill-Shannon diversity (*q* = 1, provides a balanced measure of both rare and common species), and Hill-Simpson diversity (*q* = 2, emphasizes common species) (Jost 2006). All ANOVAs included fungicide treatment, year, and fungicide treatment*year interaction as fixed effects and experimental plot as a random effect to account for repeated measures.

Significance tests were based on type 3 tests in the ‘afex’ package (Henrik Singmann et al. 2021). As a measure of the effect size for each variable in a model, we report η^2^_partial_, which is calculated as the ratio of the variance explained by the variable to the sum of that explained variance plus the residual error variance. For each variable in a model, η^2^_partial_ is bound between 0 to 1. We then evaluated differences among fungicide treatments and between years using Tukey’s Honestly Significant Difference (HSD) post-hoc tests. We checked assumptions of normality and homogeneity of variance using diagnostic plots of model residuals. To minimize heteroscedasticity, we log-transformed plant richness and biomass.

#### Plant community composition

Even if plant communities change in richness when experiencing lower disease, this may not translate to consistent shifts in the relative abundance of certain plant species that result in similar community compositions. At one end, low disease pressure could generate convergence towards a similar community state. This could happen when host species respond to disease reduction by consistently increasing or decreasing in abundance. Alternatively, communities may fail to converge in composition when stochastic colonization and loss of species occurs over time, leading to variable community structure. Thus, we also assessed whether fungicide treatments influenced plant community composition over time by modelling the relative cover of plant species using a multivariate generalized linear model in the ‘mvabund’ R package with a negative binomial distribution (Wang et al. 2012). This analysis allows us to detect both community-level and species-level responses to fungicide treatments. When considering species-level responses, we used univariate tests that were adjusted for multiple comparisons through resampling based on the Holm step-down procedure (Wang et al. 2012). We accounted for repeated sampling of communities by restricting permutations within blocks that correspond to the identity of the experimental plot using the ‘bootID’ argument (*n* = 999 permutations). Patterns of plant community dissimilarity were then visualized using nonmetric multidimensional scaling based on Bray-Curtis distances of plant relative cover in R package ‘vegan’ (Oksanen et al. 2008).

We expected the colonization and loss of plant species in our experimental plots to contribute to variation in community structure among plots over time, and we predicted that fungicide treatments would interact with this variation. Specifically, disease reduction could further increase variability in community structure by reducing selection via disease and thus increase within-treatment variation. Therefore, we also examined whether within-treatment variation in community structure, measured as the distance from the Bray-Curtis treatment centroid of each year, differed among treatments and years using a repeated measures ANOVA. We used experimental plot as a random effect to account for repeated measures, and log-transformed distance from the centroid to account for heteroscedasticity.

## Results

### Fungicide treatment effects on disease burden in tall fescue

The application of fungicide decreased cumulative foliar fungal disease burden over the growing season, as measured across the three common diseases of tall fescue (*F*_3,60_ = 176.63, *p* < 0.001, η^2^_partial_ = 0.89, Table S1, Figure S3, S4). In plots that received year-round fungicide, disease was reduced by 42.4% in 2017, 35.6% in 2018, and 85.9% in 2019 relative to the control (Table S1). In addition, disease burdens declined from 2018 to 2019 by on average 58% (*F*_2,120_ = 1610.82, p < 0.001, η^2^_partial_ = 0.96), and the effects of fungicide treatment on disease varied among years (treatment*year: *F*_6,120_ = 51.76, *p* < 0.0001, η^2^_partial_ = 0.72, Figure S3). In the first year of the experiment (2017), disease burdens from plots experiencing year-round fungicide were significantly lower than those from plots sprayed for nine months (year-round v. nine-month contrast for 2017, *p* < 0.001). In subsequent years, disease burdens between those fungicide treatment groups were not statistically distinguishable from each other (year-round v. nine-month 2018 contrast, *p*= 0.37; 2019 contrast, *p* = 0.98, Figure S3). Despite that decrease in statistical significance and the large variation among years, overall, the treatments that applied fungicide over a greater fraction of the growing season tended to reduce disease burden more over the growing season.

The community composition of foliar diseases remained relatively consistent over treatments and years. Anthracnose accounted for most of the disease in tall fescue throughout the experiment and had the highest relative contribution across all fungicide treatments and years (mean relative contribution = 0.93, range: 0.53 to 1.0). Rust (mean, = 0.11, range: 0.0 to 0.42) and brown patch (mean = 0.07, range: 0.0 to 0.39) infections were less frequent (Figure S5). Thus, although all fungicide treatments reduced disease, seasonal fungicide application did not alter parasite community composition.

### Plant diversity

Fungicide treatments generally reduced plant diversity, but with important variation among diversity metrics. Plant richness varied among years (year: *F*_2,120_ = 72.18, *p* < 0.001, η^2^_partial_ = 0.42), and was reduced by fungicide treatment (treatment: *F*_3,60_ = 3.81, *p* = 0.014, η^2^_partial_ = 0.07). Plant richness was comparable in 2017 (mean richness = 4.59) and 2019 (mean richness = 4.14; Tukey HSD *p* = 0.089), but relatively higher in 2018 (mean richness = 7.16; Tukey HSD *p* < 0.001, Figure 1, left column of panels). Still, the effect of fungicide treatment on plant richness did not change between years (treatment * year: *F*_6,120_ = 1.20, *p* = 0.32). Control plots that were never sprayed with fungicide, experiencing ambient levels of disease, on average, had 1.2 more plant species than plots treated with fungicide (Figure 1, left column of panels). Plant richness did not differ between the three fungicide-treated groups (Tukey HSD *p* > 0.05). In summary, fungicide reduced plant richness independently of the duration of fungicide exposure, and the effect of fungicide on richness was consistent across years despite changes in plant richness.

**Figure 1.**
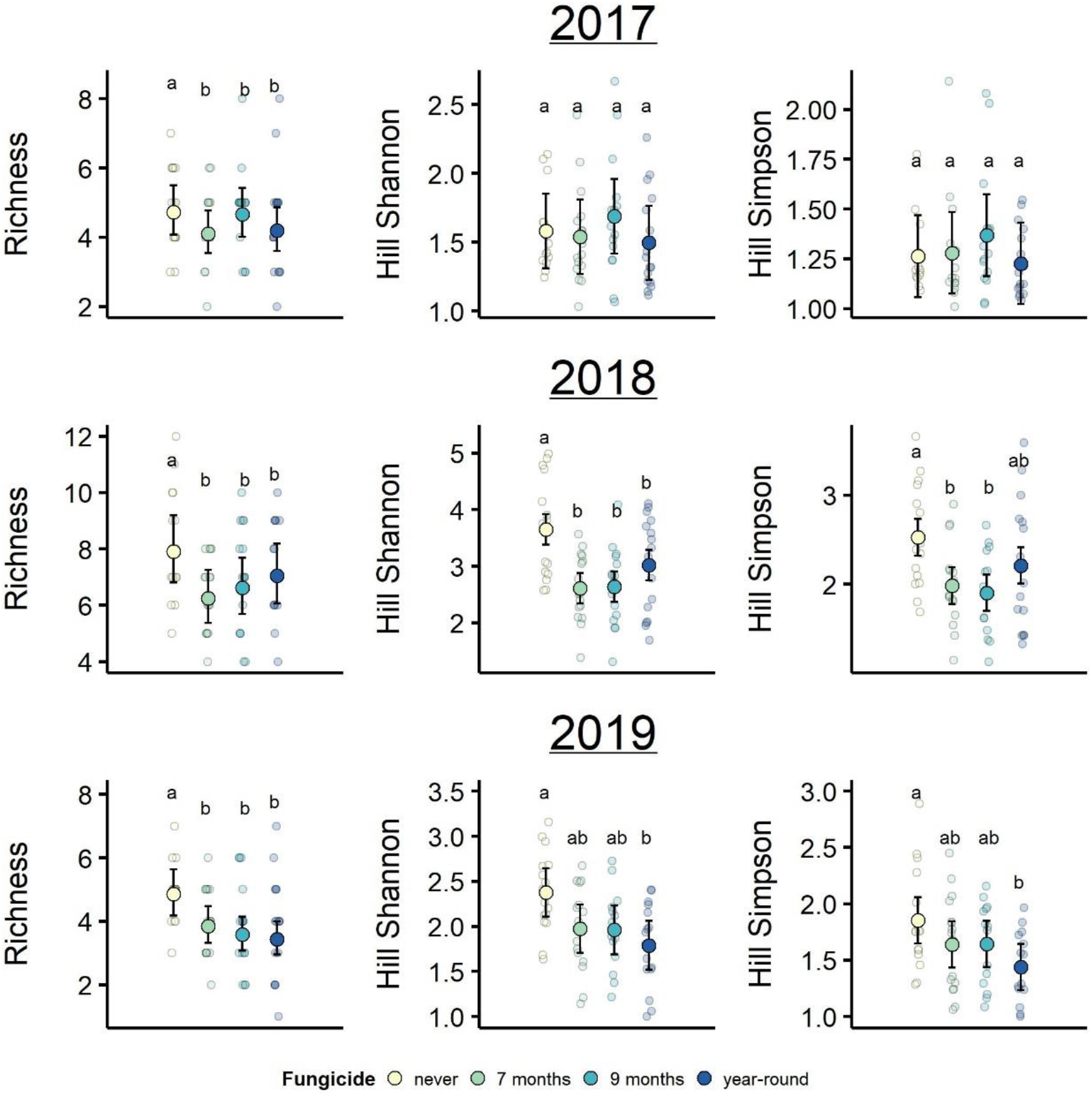
Plant diversity, measured as taxonomic richness, Hill-Shannon diversity, and Hill-Simpson diversity was generally reduced by fungicide treatments and varied across years. Means not sharing the same lower-case letters indicate significant differences between fungicide treatment groups. Due to the treatment*time interaction, Hill Shannon and Simpson diversity groupings are based only on comparisons between treatments within each year. Plotted are observed treatment means and their associated 95% confidence intervals, and smaller points display the raw data, which were jittered for visualization.

Fungicide treatment reduced plant Hill-Shannon diversity (*F*_3,60_ = 5.42, *p* = 0.002, η^2^_partial_ =0.12) and Hill-Simpson diversity, although this latter main effect was only marginally significant (*F*_3,60_ =2.68, *p* = 0.06, η^2^_partial_ =0.07). Between-year variation in both Hill-Shannon and Hill-Simpson diversity was large and a main driver of plant diversity in this experiment (year: Shannon *F*_2,120_ = 154.44, *p* < 0.001, η^2^_partial_ =0.55; Simpson *F*_2,120_ =100.85, *p* < 0.001, η^2^_partial_ = 0.44, Figure 1). Moreover, there was a significant interaction between fungicide treatment and time for both diversity measures (treatment*year: Shannon *F*_6,120_ =5.49, *p* ≤ 0.001, η^2^_partial_ = 0.12; Simpson *F*_6,120_ = 4.19, *p* = 0.001, η^2^_partial_ = 0.09). Hill-Shannon and Hill-Simpson diversity did not differ among treatment groups in 2017, the first year of experiment (Tukey HSD *p* > 0.05). In 2018, both Hill diversity metrics were higher in the control plots than the fungicide-treated plots, except for Hill-Simpson diversity in plots treated with fungicide year-round (Tukey HSD: Figure 1). Finally, in 2019 only the control and year-round fungicide-treated plots differed in plant diversity (Tukey HSD: Shannon *p* = 0.014, Simpson *p* = 0.017). In contrast to plant richness, the effects of fungicide on Hill-Shannon and Hill-Simpson diversity were more variable among fungicide treatment groups, and only became evident after the first year of the experiment.

### Plant biomass

Plant community biomass increased in response to the fungicide treatments (treatment: *F*_3,60_ = 20.27, *p* < 0.001, η^2^_partial_ = 0.25) (Figure 2). Year-round fungicide treatment increased plot biomass by 14.9% in 2017, 47.7% in 2018, and 46.6% in 2019 relative to the control (i.e., no fungicide) plots. Similar increases in plant biomass also occurred in the plots treated with fungicide for nine months (% percent change relative to control: 2017 = 12.1%, 2018 = 44.4%, 2019 = 45.3%). Increases in plant biomass from plots exposed to fungicide for seven months were about half as large as those from the other fungicide treatments (percent change relative to control: 2017 = 5.1%, 2018 = 26.7%, 2019 = 26.5%). Similar fungicide treatment effects were observed in the 2018 sorted biomass data, and differences in 2018 plant biomass were chiefly driven by an increase in fescue biomass within fungicide-treated plots (Figure S6). Across all treatments, plant biomass increased over time and was highest in 2019 (year: *F*_2,120_ = 64.14, *p* < 0.001, η^2^_partial_ = 0.42). There was weak evidence of an interaction between fungicide treatment and time on plant biomass, which may have resulted from an increase in fungicide effects on biomass in 2018 and 2019 (treatment*year: *F*_6,120_ = 1.99, *p* =0.07). Overall, differences in plant community biomass were related to variation in fungicide exposure throughout the growing season, so that communities exposed to fungicide for a longer duration had greater biomass.

**Figure 2.**
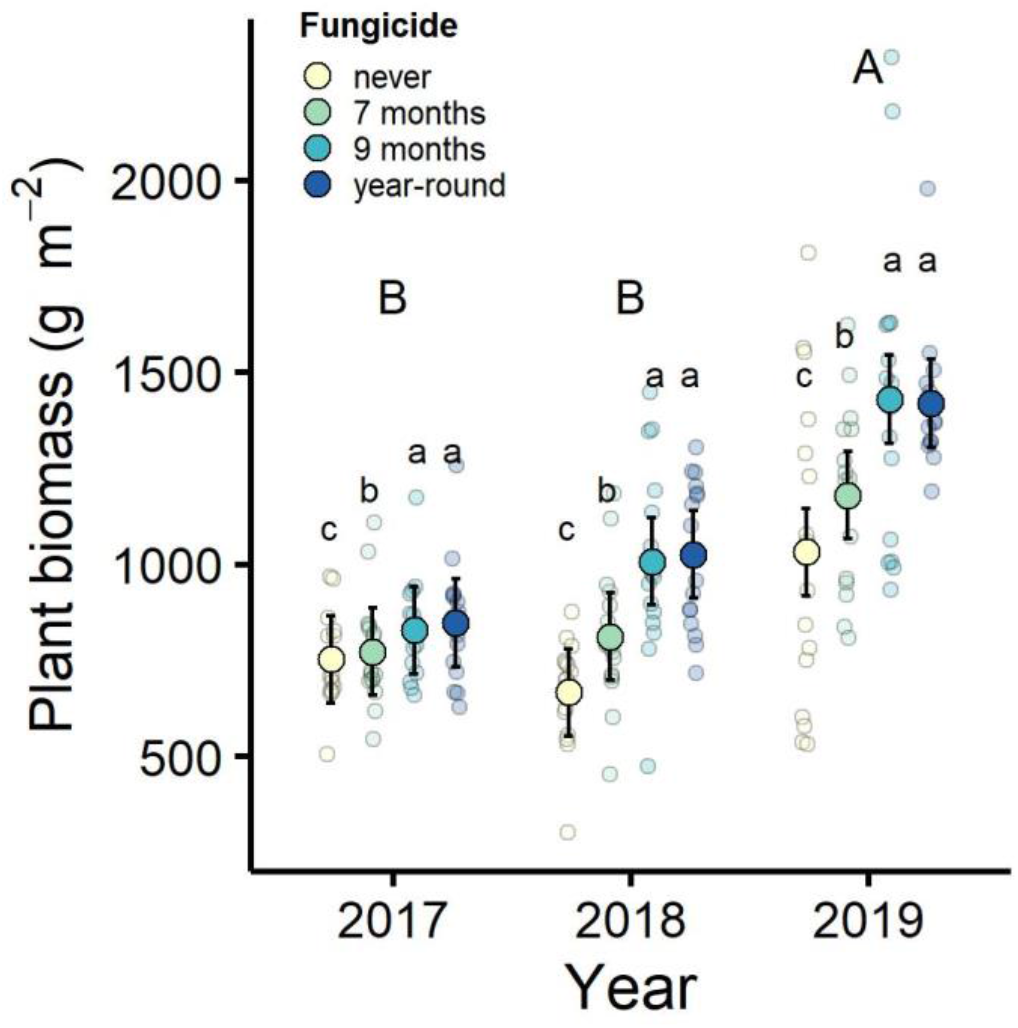
Plant community biomass generally increased in response to fungicide treatments. Years not sharing the same capital letters indicate significant differences between years based on Tukey’s HSD post-hoc tests. Treatments not sharing the same lower-case letters indicate significant differences among fungicide treatment groups. Plotted are observed treatment means and their associated 95% confidence intervals, and smaller points display the raw data, which were jittered for visualization.

### Plant community composition

Plant communities did not converge towards a similar community composition. The species-level composition of plant communities changed over time (multivariate glm: deviance= 34.73, *p* < 0.001), but was not affected by fungicide (treatment: deviance= 26.46, *p* = 0.897, Figure S7, S8). Additionally, the temporal changes in plant community composition did not interact with the fungicide treatments (treatment *time: deviance = 4.85, *p* = 0.718). For example, the relative cover of the dominant plant, tall fescue, was not affected by fungicide treatments (*p* = 0.45), but generally declined over time (univariate test, *p* > 0.001). In the first year of the experiment in 2017, all plots were dominated by tall fescue (mean relative % cover = 87.48, sd = 8.59), then fescue declined in 2018 (mean relative % cover = 62.37, sd = 16.31) and 2019 (mean relative % cover = 69.49, sd = 20.30). Similarly, tall fescue absolute percent cover was not impacted by fungicide treatments and declined over time (mean absolute % cover: 2017= 95.9, 2018 = 83.0, 2019 = 82.4), so these results were not driven by differences in total absolute percent cover of all species. We could not identify any plant species that differed in relative abundance between fungicide treatments over time (all univariate species-level tests: *p*-adjusted > 0.05). Ultimately, the fungicide treatments did not affect species-level composition of plant communities, indicating communities were not converging towards a similar community state under lower disease.

Although fungicide treatments did not affect species-level composition of plant communities, the fungicide treatments tended to increase variation in plant community composition (treatment: *F*_3,60_ = 11.05, *p* < 0.001 η^2^_partial_ =0.260). Plant community composition varied more among plots within the year-round fungicide treatment and more among plots within the nine-month fungicide treatment than within the seven-month and never-sprayed fungicide treatment groups (Tukey HSD *p* < 0.05, Figure 3). In addition, variation in plant community composition among plots increased over time (year: *F*_2,120_ = 307.01, *p* < 0.001 η^2^_partial_ =0.651), and the effect of fungicide treatments on variation in plant community composition changed over time (treatment*year *F*_6,120_ = 3.55, *p* = 0.004, η^2^_partial_ =0.061). This interaction was driven by greater increases over time in variation among the plots sprayed with fungicide for nine months, and among the plots sprayed year-round (Tukey HSD, Figure 3). This indicates that these disease-reduction treatments amplified variation in host community structure over time. While fungicide treatments did not explain differences in the average plant community composition, variation in community composition within fungicide treatment groups was greater in plots treated with fungicide for a longer time each growing season.

**Figure 3.**
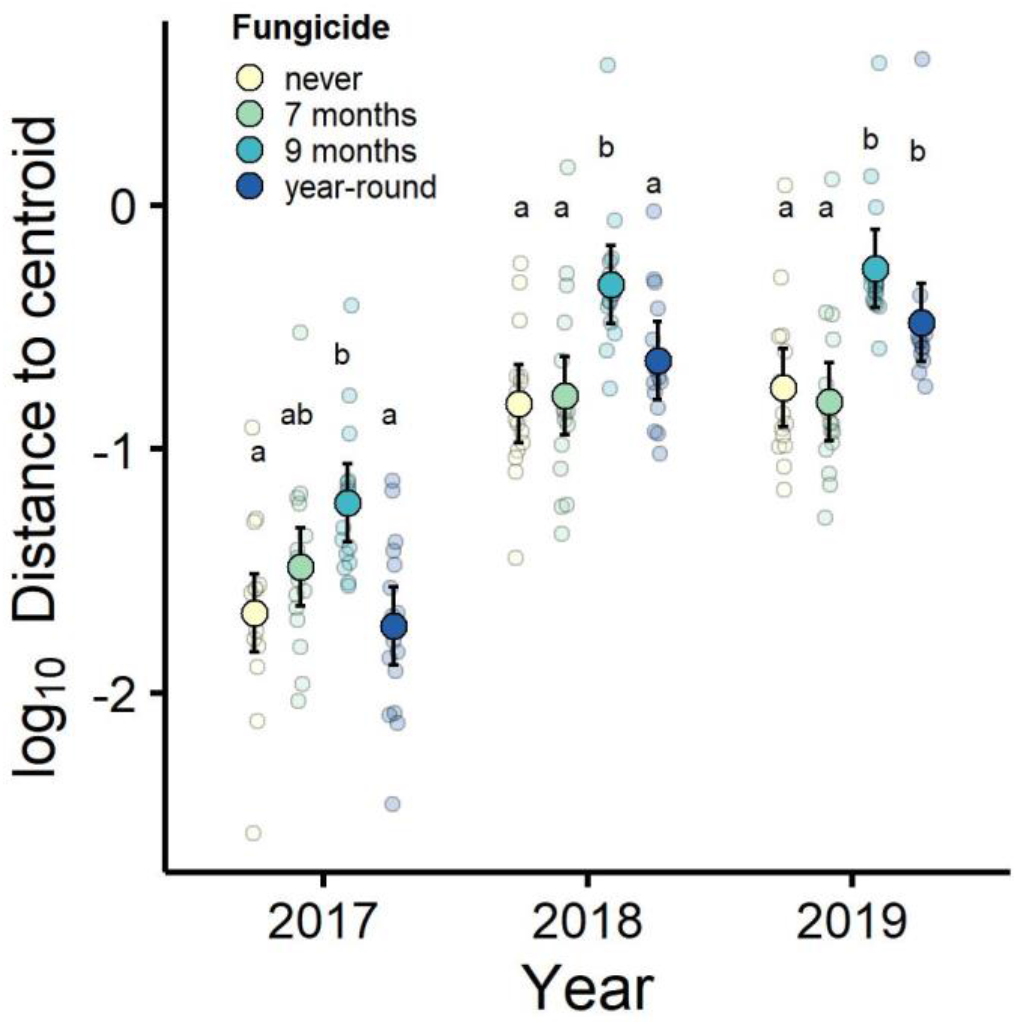
Variation in plant community composition increased over time and with seasonal duration of fungicide treatment. The fungicide treatments tended to increase variation in plant community composition, and that increase in variation tended to amplify over time, particularly in plant communities treated with fungicide for nine months out of the year or year-round. Within-treatment variation in plant community composition was measured within years as the log-transformed distance to the Bray-Curtis community centroid. Plotted are observed treatment means on log-transformed data and their associated 95% confidence intervals, and smaller points display all data, which were jittered for visualization. Treatment means not sharing the same letters denote significant differences based on Tukey’s post-hoc comparisons of fungicide treatments within each year.

## Discussion

Foliar fungal infections are ubiquitous, and yet their effects on plant community structure among communities and over time are not well known. Here, experimentally reducing foliar fungal disease in intact plant communities generally decreased plant diversity and increased plant biomass. However, disease reduction did not lead to consistent shifts in plant composition and communities did not converge towards a similar composition under lower disease. Instead, plant community composition varied considerably among experimental plots and over time, and this among-plot variation in plant composition was amplified by disease reduction. Overall, our three-year experiment suggests that foliar fungal diseases can not only maintain or change plant diversity, but can modulate the spatiotemporal dynamics of plant community composition.

Variation in disease did not explain differences in average plant community composition among plots (indicating no community convergence within treatment groups), but played a notable role in reducing variation among communities. Consistent shifts in plant species composition and abundance in response to disease reduction did not occur in our field experiment. Consequently, community assembly did not appear to be deterministic. Among our plots, stochastic colonization and extinction of plant species occurred, as evidenced by the high variation in community composition among plots throughout the experiment. Thus, although plant communities responded to the fungicide treatments, as shown by richness and Hill-diversity metrics, plant species did not respond in a consistent way that would result in convergence of plant communities towards a similar community state within a treatment group. Disease reduction may not favor specific plant species in this system, or we could not detect species-level selection given the large heterogeneity in community composition. Plant community composition was variable among experimental plots and this variation could be related to dispersal limitation in our system (Martin and Wilsey 2012, Collins et al. 2017) along with a tendency for communities to exhibit ecological drift. In contrast to any signals of community convergence in response to disease exclusion, plant community composition was more variable among plots that were treated with the fungicide. Being released from disease appears to amplify variation in community structure in our system. Weaker selection can heighten the importance of other processes that govern community structure, so being released from disease may have made communities more influenced by stochastic processes. Herbivore and predator exclusion experiments have reported similar findings to our study and may point to a broader role of natural enemies in reducing variation in community composition by reducing the relative contribution of stochastic processes (Chase et al. 2009, Mortensen et al. 2018, Chen et al. 2022).

Over the three years of this experiment, plant communities changed considerably, starting as primarily dominated by tall fescue and becoming relatively more diverse. These shifts in plant communities may have resulted in part from a shift in the timing of annual mowing: the experiment was mowed the summer before the experiment was implemented (i.e., in 2016), and thereafter, was mowed during the late fall. Mowing later in the year may have allowed species other than tall fescue, particularly dicots, to grow undisturbed through the summer and fall. This could potentially increase plant establishment from seed, survivorship, growth from root stocks, and size at the end of the growing season, all of which could have led to increased plant diversity over time. Despite changes in plant communities over time, disease reduction consistently lowered plant richness in our experiment. In contrast, Hill-Shannon and Hill-Simpson diversity metrics, which place weight on species abundance, revealed treatment effects that varied over time. In the first year of the experiment, Hill-Shannon and Hill-Simpson diversity were similar among fungicide treatment groups, and this was likely due to the dominance of tall fescue among all plots. This result reflects the high initial dominance of tall fescue among all plots. It may also reflect an underestimation of plant diversity in the first year, when the plant community survey was performed a month later (November) than in subsequent years (October). By November, some species may have senesced and not been detected. In subsequent years, after tall fescue declined in both absolute and relative abundance across plots, Hill-Shannon and Simpson did differ among fungicide treatments and diversity was generally lower in fungicide-treated plots. Foliar fungal disease may increase (as shown here), decrease, or have no effect on plant diversity (Peters and Shaw 1996, Mitchell 2003, Allan et al. 2010). These effects of disease on host diversity may interact with host temporal dynamics and further alter the trajectory of host community assembly (Kardol et al. 2006, Jiang et al. 2020, Szefer et al. 2020, Wilfahrt et al. 2020, Heckman et al. 2022). In our system, the initial decline of the dominant grass, tall fescue, along with the colonization and extinction of other plants that were potentially released from disease may have resulted in the observed effects on plant diversity over time.

Plants are exposed to a diverse array of parasites that impact their fitness (Mordecai 2011, Bever et al. 2015), so plant diversity may further be affected by changes in the parasite composition (Cappelli et al. 2020). Variation in fungicide impacts could occur via additive effects of disease burdens (Dantec et al. 2015) and/or shifts in parasite composition (Cappelli et al. 2020). In addition, differences in parasite community structure can alter within-host interactions that scale up to impact disease epidemics (Halliday et al. 2017, Clay et al. 2020). In our experiment, treating plant communities with fungicide for a longer time generally reduced disease more. However, differences in the overall disease burden among plots in different fungicide treatments did not result in further differences in plant richness among treatment groups. Despite timing the fungicide treatments to fit the phenology of the foliar fungal parasites in our system, the treatments had little effect on parasite community composition, so our experiment was not able to test effects of parasite community composition on host diversity. While we could not test effects of parasite community composition, the reduction of plant diversity under fungicide treatment, with relatively little variation in effects among the three fungicide treatments, suggests that the effects of fungicide treatment on plant diversity were mediated by something shared among fungicide treatments. For example, in all treatments, fungicide reduced disease in spring and early summer, key seasons for growth of tall fescue.

Interactions between plant communities and disease are reciprocal in nature; disease plays a role in mediating host diversity (Minchella and Scott 1991, Allan et al. 2010, Mordecai 2011) and at the same time disease transmission and diversity are also driven by host community structure (Halliday et al. 2019, 2021). These feedbacks between disease and host communities may occur on different timescales (Halliday et al. 2019). For example, in our system, disease diversity and abundance tend to peak later in the growing season (Halliday et al. 2017, Figure S2), while some disease impacts on host communities (e.g., richness effects) appear to occur earlier, in spring and early summer, as indicated by the similar effects of the different fungicide treatments on plant richness. Therefore, differences in disease burden/abundance may not be directly related to disease impacts on host communities. Scenarios like this may occur if disease diversity or abundance peaks later or earlier than critical time points of host growth or colonization, which help shape communities. Moving forward, incorporating more temporal components, such as varying fungicide treatments within a growing season, into studies on the feedback between disease and communities (Halliday et al. 2019) may provide further insight into these complex dynamics.

Along with its impacts on host diversity, disease can reduce plant biomass and contribute to variation in ecosystem productivity (Mitchell 2003, Allan et al. 2010, Seabloom et al. 2017). Here, exposure to fungicide and consequently lower levels of disease throughout the growing season generally led to greater plant community biomass. In addition, a longer duration of fungicide exposure further increased plant biomass. Overall, these impacts of disease on plant community biomass were strong and resulted in a ∼50% decline in productivity in certain years. Such prominent changes in biomass and primary productivity could have further implications for other consumers in our system. The strong response of plant biomass to differences in disease in our field experiment provides another important example of the effects of diseases on ecosystems (Mitchell 2003, Seabloom et al. 2017, Zaret et al. 2022).

Similar to predators and other natural enemies (Chase et al. 2009, Chen et al. 2022), parasites interact with their hosts in ways that may shape the relative importance of deterministic and stochastic processes throughout community assembly. While disease reduction did not result in one signature of deterministic community assembly, plant communities that converged in composition, disease did reduce variation in community structure. When disease was reduced, communities tended to become less diverse and more variable, suggesting that the relative importance of stochastic processes may increase under lower disease (i.e., under weaker selection). Given that disease could be an agent of selection in most systems, resolving its role in either reducing or amplifying variation among communities may yield insights into its consequences for the trajectory of community assembly. Taken together, our study suggests that while disease impacts on host communities may be chiefly driven by selection in conjunction with other deterministic processes like competition, how disease shapes communities over time also can depend on stochastic processes structuring those communities.

## Data Availability Statement

The data and code that support the findings of this study are available through Zenodo at https://zenodo.org/badge/latestdoi/283863948

## Acknowledgments

For their assistance with field work, we thank: Anita Simha, Brandon Wheeler, Matthew Carey, Storm Crews, Charles Muirhead, Jordan Link, Claire Thefaine, Safiyyah Motaib, Julia Knorr, and Ryan Cook. We thank Peter Morin and members of the Morin lab for their feedback. This work was supported by the NSF-USDA joint program in Ecology and Evolution of Infectious Diseases (USDA-NIFA AFRI grant 2016-67013-25762). The findings and conclusions of this publication are those of the authors and should not be construed to represent any official USDA or U.S. Government determination or policy.

## Author contributions

RLG analyzed the data and wrote the first draft of the manuscript. FWH and CEM conceptualized the study. FWH and KRO designed the experiment. FWH, KRO, BNJ, and RWH collected the data. All authors contributed to revisions of the manuscript.

