## Supplemental materials for "Disease decreases variation in host community structure in an old-field grassland"

**Supporting information for:** Disease decreases variation in host community structure in an old-field grassland

| Year | Treatment | % change | LRR | 95% CI | |
|---|---|---|---|---|---|
| | 7 months | -12.5 | -0.13 | (-0.15, | -0.11) |
| 2017 | 9 months | -28.5 | -0.34 | (-0.40, | -0.27) |
| | year-round | -42.4 | -0.55 | (-0.62, | -0.49) |
| | 7 months | -13.9 | -0.15 | (-0.18, | -0.12) |
| 2018 | 9 months | -27.9 | -0.33 | (-0.42, | -0.23) |
| | year-round | -35.6 | -0.44 | (-0.52, | -0.36) |
| | 7 months | -63.3 | -1.00 | (-1.16, | -0.85) |
| 2019 | 9 months | -82.5 | -1.74 | (-2.03, | -1.44) |
| | year-round | -85.9 | -1.96 | (-2.22, | -1.72) |

Table S1. Log response ratios (LRR) of fescue disease AUDPS in response to fungicide treatments. Reported are % change in AUDPS relative to the control, bootstrapped LRR and 95% confidence intervals.

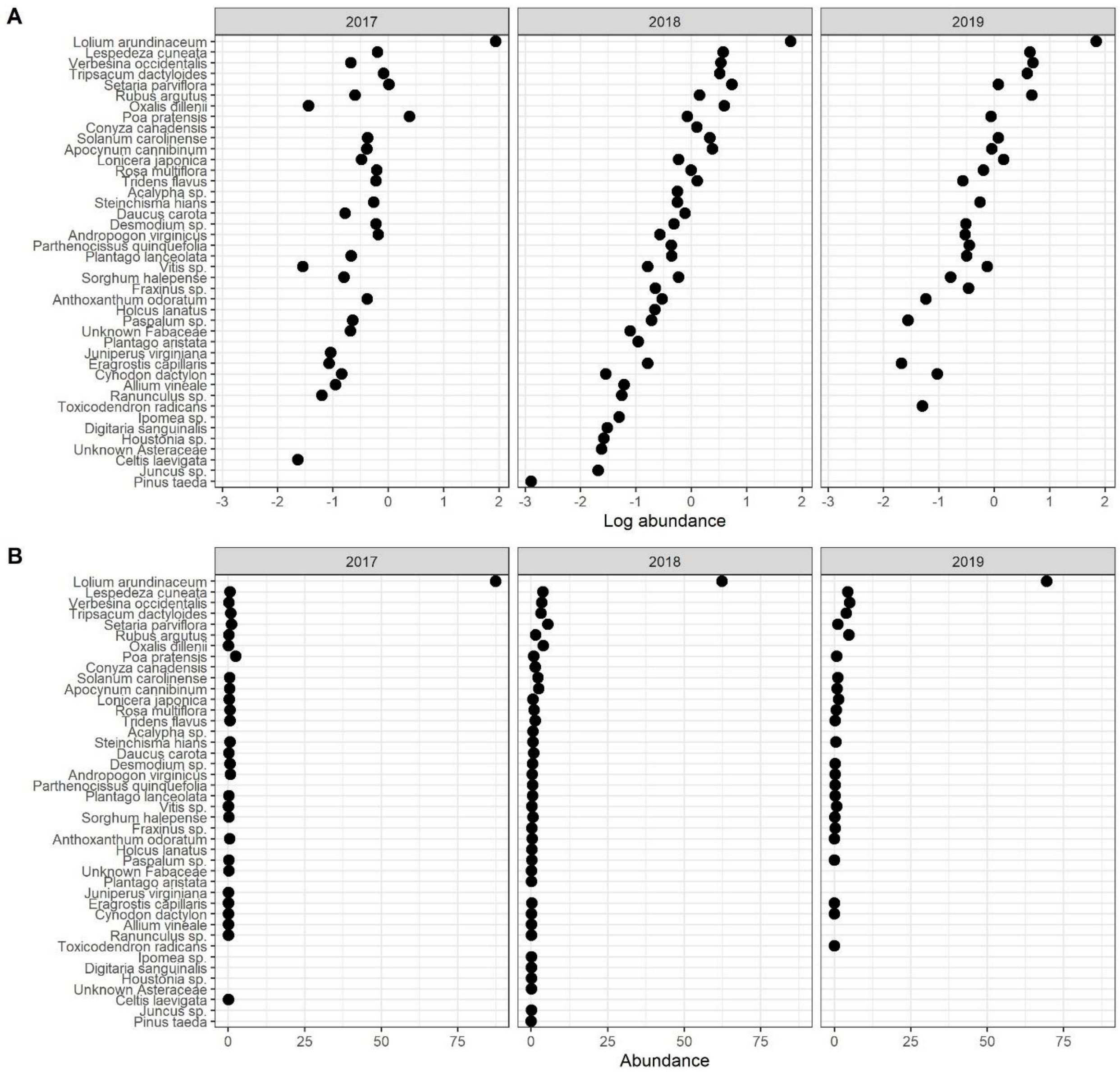


**Figure S1.** Ranked log transformed abundance (A) and raw-untransformed abundance (B) of all plant species across years in experimental plots. Relative abundance was averaged across all 64 experimental plots for a given year. Tall fescue (*Lolium arundinaceum)* was the dominant plant species each year.

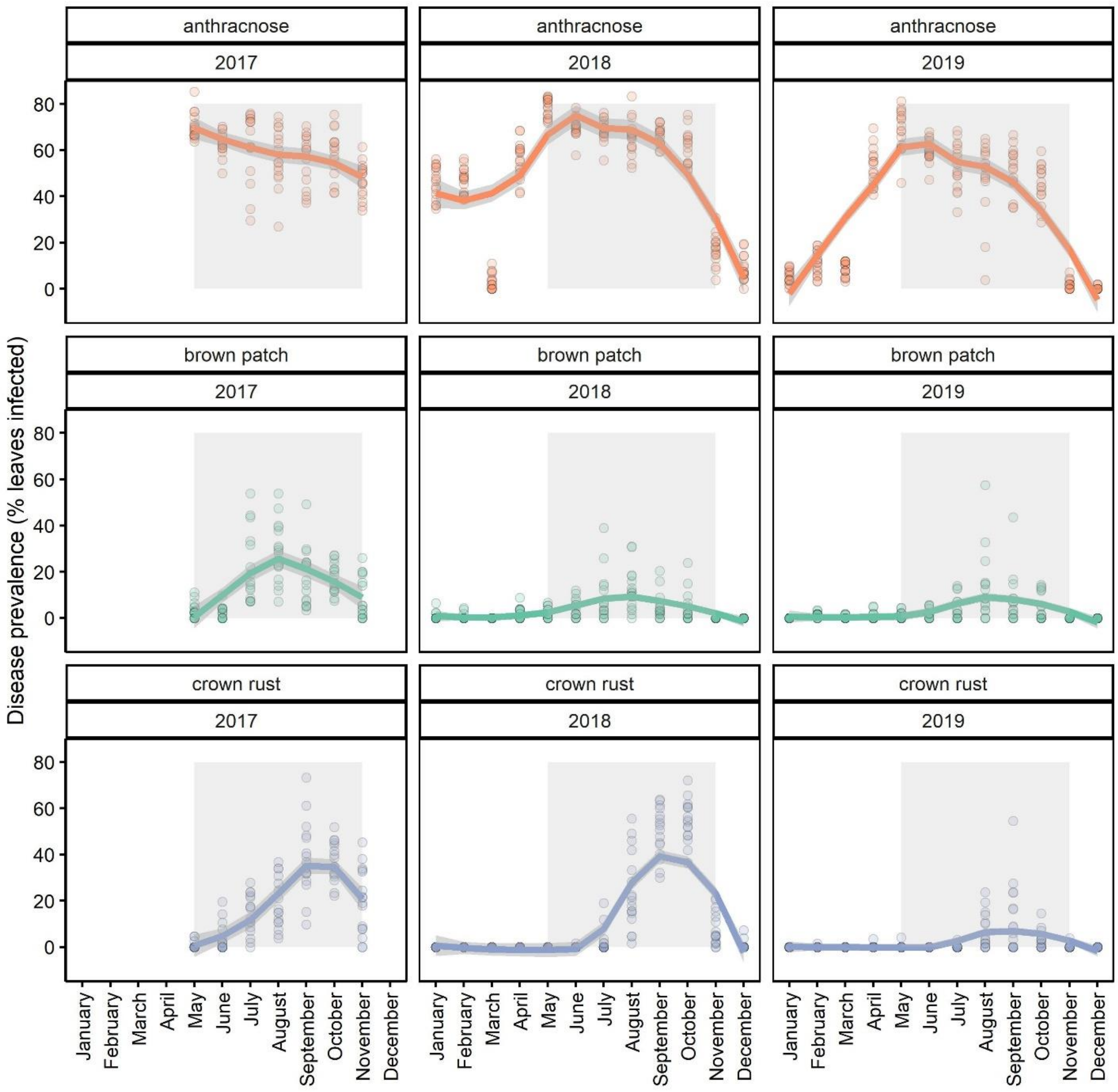


**Figure S2** Seasonal disease epidemics on leaves of tall fescue from control (never sprayed with fungicide) plots. Generally, anthracnose peaked in late spring, while brown patch peaked in summer and crown rust in fall. The grey rectangle delineates the disease survey data used to calculate disease burden in analyses (i.e., AUDPS of disease prevalence from May-November). Loess fitted lines are drawn to show the peak prevalence of each disease, and points are the raw disease prevalence data for each of the 16 control plots.

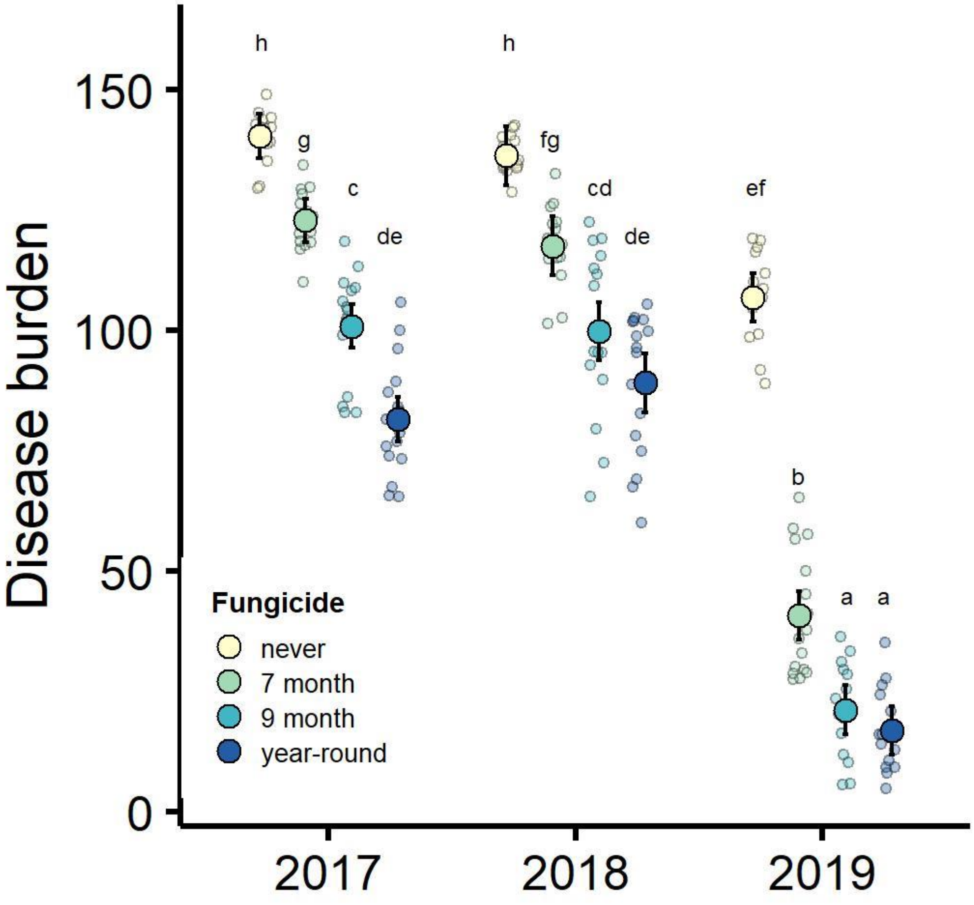


**Figure S3.** Disease of tall fescue, quantified as the cross-species disease burden (AUDPS) based on prevalences of the three focal diseases (anthracnose, brown patch, and crown rust), was strongly reduced by fungicide treatments and these treatment effects varied across years. Plotted are observed treatment means and their 95% confidence intervals, and smaller points represent the raw data that are jittered to show the distribution of the data. Lower case letters denote groups based on Tukey post-hoc tests using all pairwise comparisons to compare treatment effects over time.

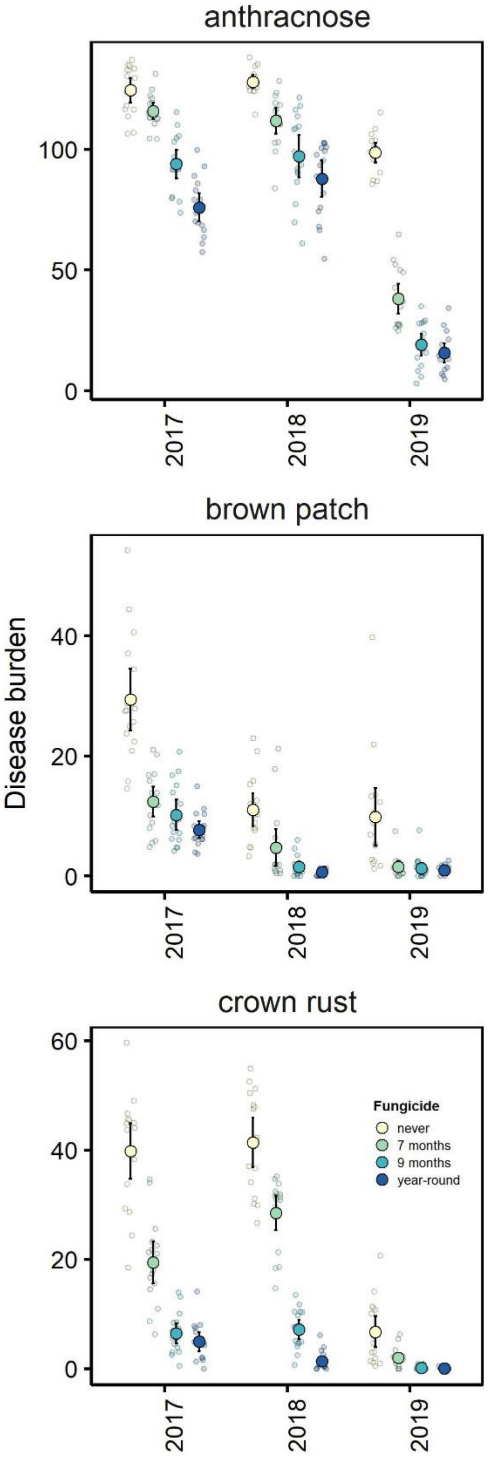


**Figure S4.** The effects of fungicide treatments on disease burden (AUDPS) of three diseases of tall fescue (anthracnose, brown patch, crown rust). Large points indicate treatment means with their 95% confidence intervals, while smaller points are the raw data. Raw data points are jittered to show the distribution of the data.

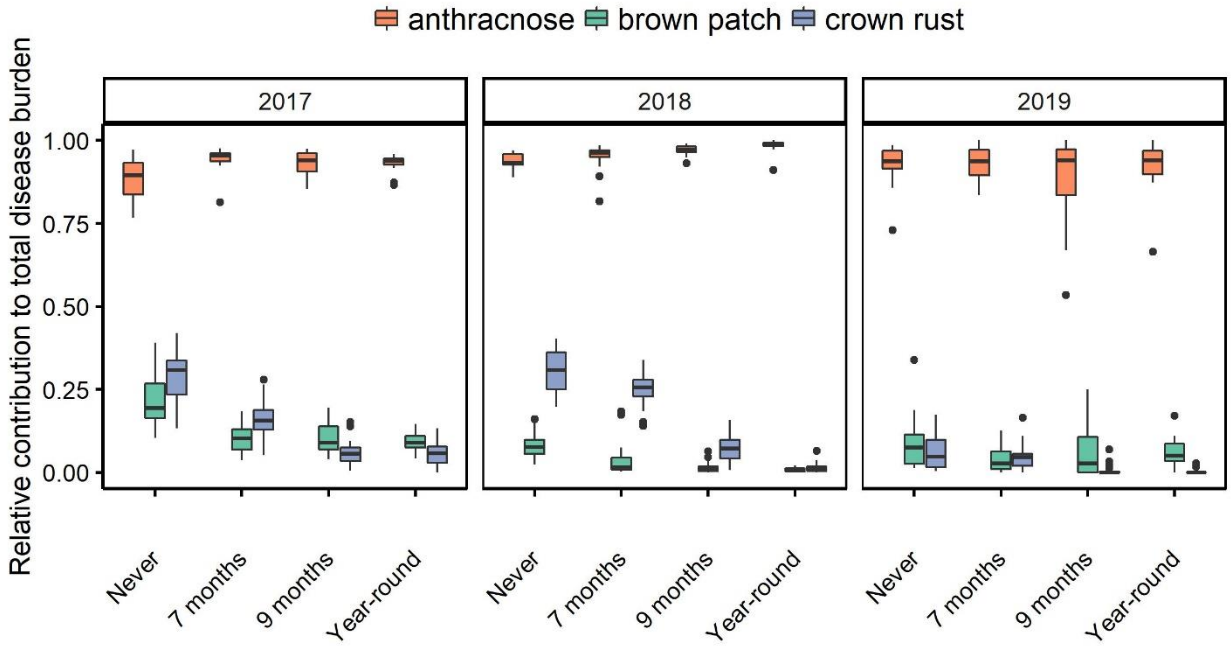


**Figure S5**. The relative contribution of anthracnose, brown patch, and crown rust to the overall cross-disease disease burden (AUDPS). The disease anthracnose was the chief contributor to total disease across all treatments and years.

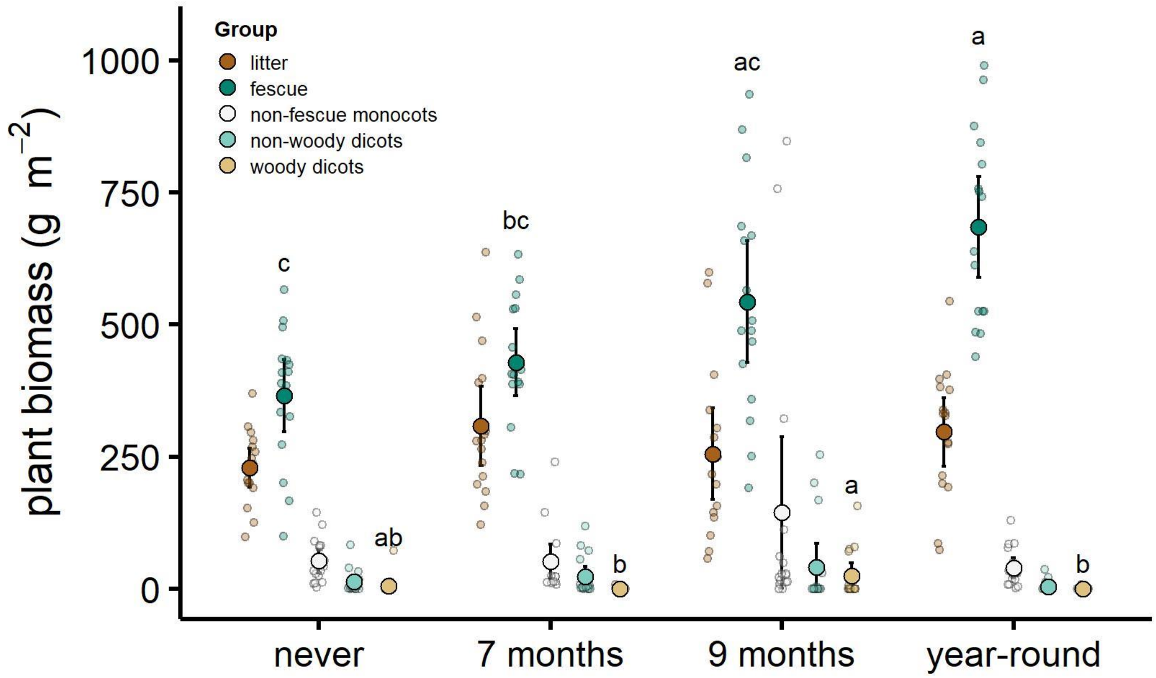


**Figure S6.** In 2018, the aboveground biomass of certain plant groups differed among fungicide treatments (MANOVA, treatment: $F_{3,60} = 4.23$, Wilks $\Lambda = 0.20$, $p < 0.0001$). Specifically, differences among treatments in 2018 plant biomass were attributed to tall fescue (ANOVA, treatment: $F_{3,60} = 11.47$, $p < 0.0001$, $\eta^2 = 0.36$) and woody dicots ($F_{3,60} = 3.32$, $p = 0.03$, $\eta^2 = 0.14$). Litter ($F_{3,60} = 1.32$, $p = 0.28$), non-fescue monocots ($F_{3,60} = 1.87$, $p = 0.14$), and non-woody dicots ($F_{3,60} = 1.72$, $p = 0.17$) were not affected by fungicide treatments in 2018. Letters denote post hoc comparisons between treatments within each plant biomass group. Plotted are observed treatment means and their associated 95% CI, and smaller points represent the raw data that are jittered to show the distribution of the data.

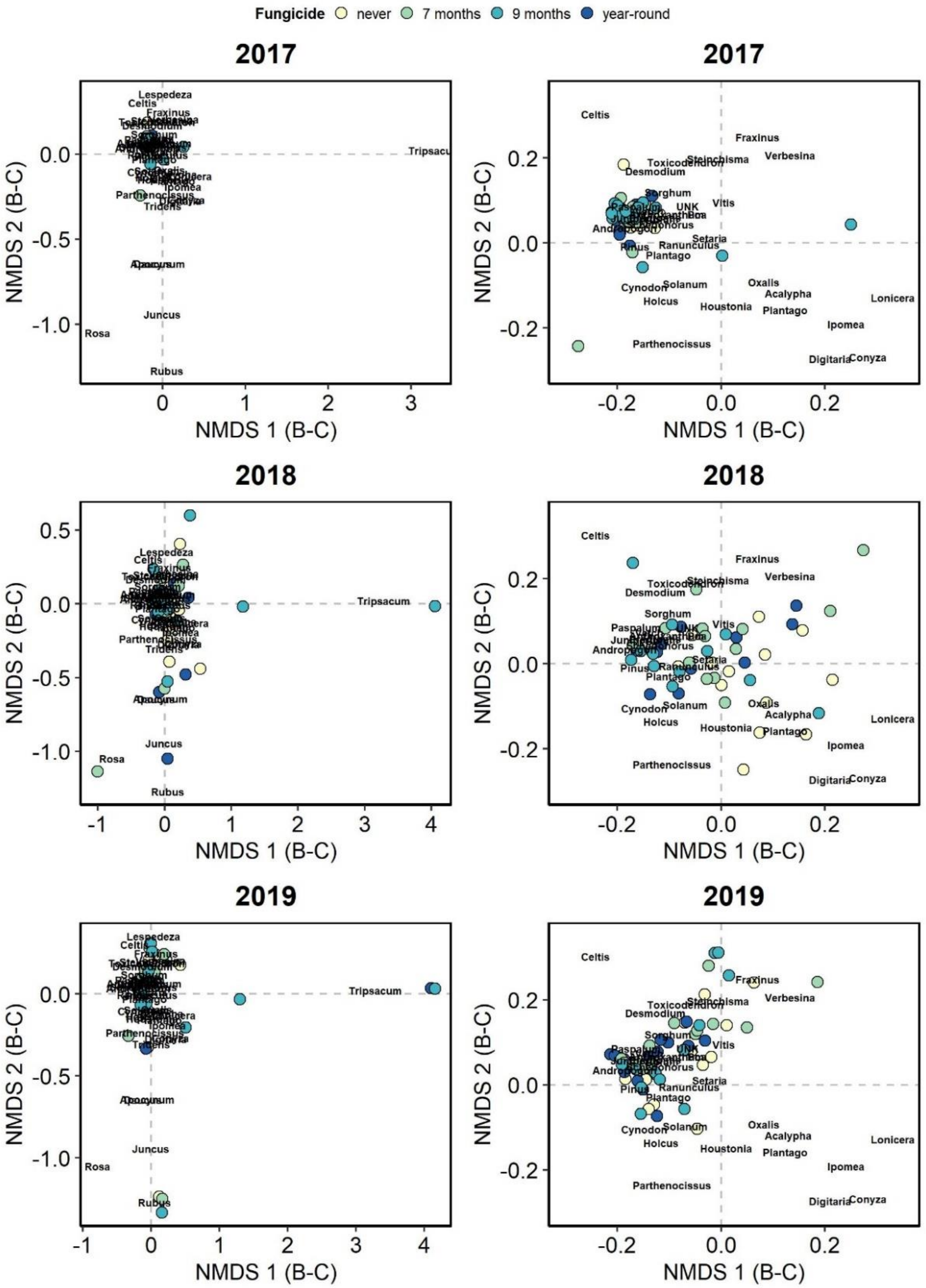


**Figure S7.** Plant community dissimilarity patterns varied across years, and were not affected by fungicide treatment. Shown are results of nonmetric multidimensional scaling of plant communities based on Bray-Curtis distances. All data are plotted on the same axes for A, C, E to show all community dissimilarity patterns. Panels B, D, F, are zoomed in to show the core community patterns. Species vectors show the distribution of plant taxa among communities. NMDS Stress = 0.12.

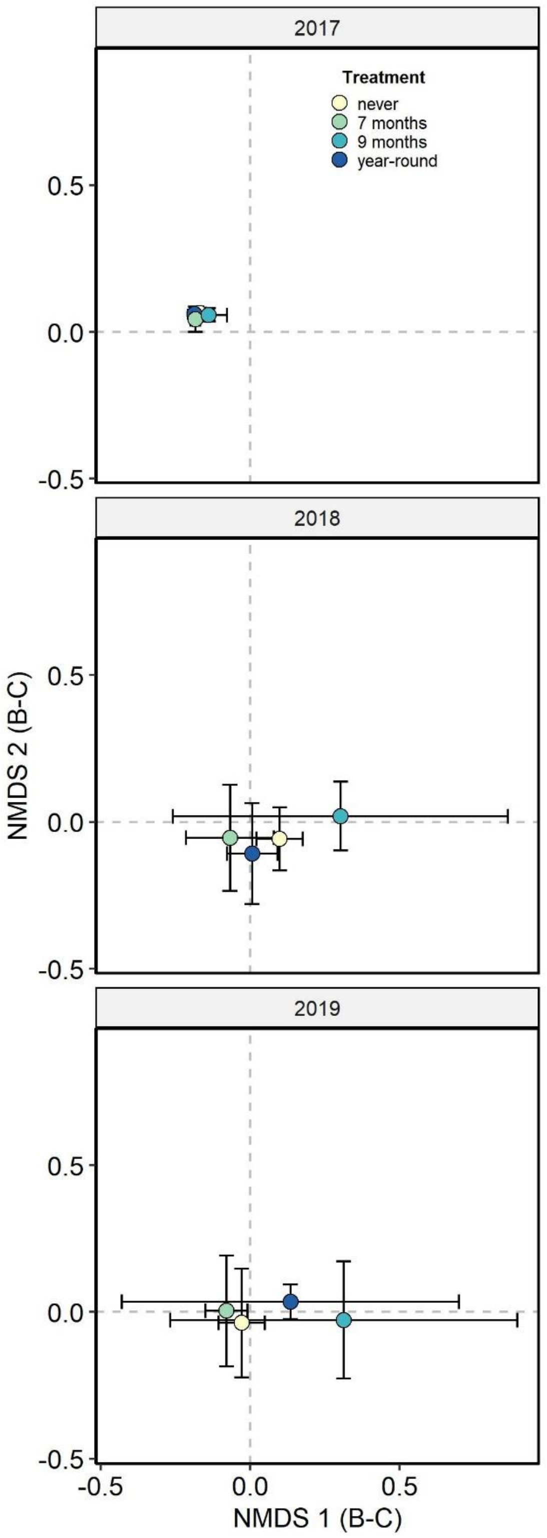


**Figure S8.** Plant community composition varied across years and was not impacted by fungicide treatments. Plotted are treatment centroids of nonmetric multidimensional scaling based on Bray-Curtis distances and their 95% confidence intervals. NMDS Stress = 0.12.
